# Unpacking Unstructured Data: A Pilot Study on Extracting Insights from Neuropathological Reports of Parkinson’s Disease Patients using Large Language Models

**DOI:** 10.1101/2023.09.12.557252

**Authors:** Oleg Stroganov, Amber Schedlbauer, Emily Lorenzen, Alex Kadhim, Anna Lobanova, David A. Lewis, Jill R. Glausier

## Abstract

**Objective:** The aim of this study was to make unstructured neuropathological data, located in the NeuroBioBank (NBB), follow FAIR principles, and investigate the potential of Large Language Models (LLMs) in wrangling unstructured neuropathological reports. By making the currently inconsistent and disparate data findable, our overarching goal was to enhance research output and speed.

**Materials and Methods:** The NBB catalog currently includes information from medical records, interview results, and neuropathological reports. These reports contain crucial information necessary for conducting in-depth analysis of NBB data but have multiple formats that vary across different NBB biorepositories and change over time. In this study we focused on a subset of 822 donors with Parkinson’s disease (PD) from 7 NBB biorepositories. We developed a data model with combined Brain Region and Pathological Findings data at its core. This approach made it easier to build an extraction pipeline and was flexible enough to convert resulting data to Common Data Elements (CDEs), a standardized data collection tools used by the neuroscience community to improve consistency and facilitate data sharing across studies.

**Results:** This pilot study demonstrated the potential of LLMs in structuring unstructured neuropathological reports of PD patients available in the NBB. The pipeline enabled successful extraction of detailed tissue-level (microscopic) and gross anatomical (macroscopic) observations, along with staging information from pathology reports, with extraction quality comparable to manual curation results. To our knowledge, this is the first attempt to automatically standardize neuropathological information at this scale. The collected data has the potential to serve as a valuable resource for PD researchers, facilitating integration with clinical information and genetic data (such as genome-wide genotyping and whole genome sequencing) available through the NBB, hereby enabling a more comprehensive understanding of the disease.

## INTRODUCTION

Effective data modeling of biological experiment data can have a major impact on downstream data usage, accessibility, and significantly improve research output. Having a robust and sound data model allows FAIR (Findability, Accessibility, Interoperability, and Reusability) data principles to be employed and provide major benefits to research progress [1]. A recent cost-benefit analysis by the European commission on FAIR data suggests that not using FAIR data principles costs the European economy approximately €10.2 billion per year [2]. Thus, improving the quality of data models by applying these principles serves to save a considerable amount of time and resources, further advancing research efforts.

With the advent of high-performance computing and artificial intelligence (AI), technologies such as natural language processing (NLP) and large language models (LLMs) can be used to facilitate data FAIRification of unstructured data [3,4]. LLMs are advanced AI systems trained on vast amounts of text data, capable of understanding and generating human-like text. These models, such as OpenAI’s Generative Pre-trained Transformers (GPT) [5] use deep learning techniques to learn from the statistical associations between words in large online text corpora, which allows them to produce human-like text outputs [4]. In the context of biomedical research, LLMs are currently being explored as a means to extract data, identify patterns, and uncover insights that may have been previously hidden [6,7]. While the current gold-standard of data curation is to perform manual curation, this process is time intensive and can introduce errors. Manual curation can also comply with FAIR principles when properly implemented. However, LLMs offer potential advantages such as increased speed, scalability, and consistency in data extraction across large datasets. It’s important to note that automated curation methods, including those using LLMs, can also introduce errors. However, LLMs may hold value in accelerating data curation and allowing FAIR data principles to be applied, ultimately improving research efficiency.

The NIH-funded NeuroBioBank (NBB, https://neurobiobank.nih.gov/) was established in September 2013 as a national resource for investigators utilizing post-mortem human brain tissue and related biospecimens for their research to understand conditions of the nervous system such as Alzheimer’s Disease (AD), Parkinson’s disease (PD), frontotemporal dementia (FTD), and many others. The NBB operates through several biorepositories across the United States, each maintaining collections of human post-mortem brain tissue and related biospecimens. These biorepositories serve as the primary sources for the specimens and associated data in the NBB catalog. The overall goals of the NBB are to 1) increase the availability of brain tissue from individuals affected and unaffected by brain disorders, 2) facilitate brain tissue distribution and 3) provide a central resource of best practices and protocols to the research community. Comprised of medical records, interview results, and neuropathological reports, the NBB catalog is an invaluable source of data for researchers. The catalog has information on clinical diagnosis, medical history, as well as results of whole genome sequencing. However, key pieces of data – specifically, the results of gross and microscopic examination of brain samples exist primarily as unstructured notes, often in the form of PDF pathological reports. The lack of standardization and inconsistent formats used across NBB biorepositories presents a significant data accessibility challenge, which hinders effective data usability and ultimately, research output. Converting these reports to a standardized format in accordance with FAIR principles would avoid duplicating efforts by different groups to extract data, accelerating research progress. Specifically, our study addresses the following FAIR principles:

- Findability: By converting unstructured reports into a standardized format, we enhance the discoverability of the data.
- Accessibility: Standardization improves data retrieval and access for researchers.
- Interoperability: A common format allows for easier integration with other datasets and systems.
- Reusability: Structured, standardized data is more readily reusable for various research purposes.

Standard representation of pathology data would provide researchers with powerful tools to understand the mechanisms underlying the development of various pathologies, ultimately leading to improvements in the diagnostics and treatment of these debilitating conditions.

In this study, we investigated the potential of LLMs in unpacking unstructured neuropathological reports, with a focus on a subset of patients with PD. Parkinson’s disease was selected for this study for several reasons. First, it provided a sizeable yet manageable dataset of 822 reports, which is large enough to be representative but not so vast as to overwhelm initial pipeline development. Second, PD exhibits moderately homogeneous pathological features, which is advantageous for developing and validating our extraction methods, as it allows for more consistent patterns in the reports while still presenting some variability. This level of homogeneity is not always present in other neurological disorders, which might have more diverse or complex pathological presentations. Additionally, PD is a prevalent neurodegenerative disorder affecting millions worldwide, making this research clinically relevant and potentially impactful.

Neuropathological reports spanned seven different biorepositories and utilized 15 different formats, including variations in file types (e.g., pdf, docx, xlsx) and report structures (e.g., narrative descriptions, region-based groupings, electronic data capture systems). Reports were first preprocessed and converted to HTML file format, as they were provided in various formats. Information was then extracted from parsed reports using a questionnaire-based method that employed few-shot learning using the *gpt-3.5-turbo* model [8]. The extracted data was then combined, harmonized, and manually reviewed for accuracy.

## MATERIALS AND METHODS

### Source data

We used 822 PD reports generated from seven NBB sites: University of Maryland, University of Pittsburgh , National Institute of Mental Health (NIMH), University of Miami, Sepulveda, Harvard Brain Tissue Resource Center, and Mount Sinai/Bronx VA Medical Center. This represents approximately 5% of the total number of reports collected by the NBB. The sites provided reports in various file formats such as pdf, docx, or xlsx.

Most of the reports contained sections outlining the specimen received, neuropathological diagnosis, macroscopic and microscopic pathological findings, and pathologist comments. Details and formats of the reports differed among the sites: for some sites, such as Maryland and Sepulveda, pathology descriptions were provided as narratives with sequential descriptions of findings. Other sites such as Harvard and Miami, grouped findings by brain regions. Notably, the Mount Sinai site utilized an electronic system to capture information, significantly streamlining the data collection process. In total, we compiled data from 822 neuropathological reports, spanning a 32-year period from 1990 to 2022. Report selection was based on the presence of a PD clinical diagnosis. No additional stratification based on age, gender, or disease stage was made, and all personally identifiable information was redacted by NBB staff. To ensure privacy and confidentiality, the NBB implemented a thorough anonymization process before sharing the data. This process included removing names, dates of birth, addresses, and any other identifiers that could potentially link the data to specific individuals. The research team received and analyzed only de-identified data, and no attempts were made to re-identify individuals. All data handling and storage procedures were in compliance with relevant data protection regulations and institutional ethical guidelines."

To facilitate the development of an automatic extraction pipeline, we classified reports based on their formats and created a subset of 65 reports, which included the most representative reports for each site and format. This selection process prioritized capturing the diversity of report formats across different sites to develop a robust and generalizable NLP pipeline, rather than aiming for a strictly random sample. The number of reports per format and site varied, reflecting the uneven distribution of PD reports across sites in our dataset. Information on the composition subset of reports for manual curation set is provided in the supplementary tables S3 and S4. The 65 reports were manually curated by two PhD-level annotators with neuroscience experience. The results were regularly discussed and reviewed by the NBB working group to ensure consistency and accuracy. This curated subset served as a "gold standard" to develop and improve the NLP pipeline.

#### Data preprocessing and parsing

Neuropathological reports in pdf file format were converted to HTML using the ABBYY FineReader Optical Character Recognition (OCR) tool (https://pdf.abbyy.com/). Similarly, docx reports were converted to HTML using the doc2html Python library (https://github.com/chadwickcole/doc2html). HTML was chosen as the target file format because it preserved information about text styles, which was utilized to mark the beginning of report sections. During preprocessing, the reports were split into sections such as gross pathology, microscopic findings, diagnosis, and sample information. For the reports where tissue information was available, an additional sub-rubric "tissue" was added. An output table was created containing report ID, section type, and section content. The data preprocessing and parsing stage enabled the conversion and organization of neuropathological reports into a structured and machine-readable format, laying the foundation for subsequent NLP pipeline development and data extraction.

### Data model development

The data model is the foundation for our approach to standardizing and structuring neuropathology reports. By data model, we refer to a representation of entities, their relationships, and allowed attributes. To design the data model, we focused on establishing a "region-finding-qualifier" triad (for example “Hippocampus – Neurofibrillary degeneration – Moderate”). This approach contrasts with many Common Data Elements (CDEs), where predefined combination of regions and finding are treated as fixed, single data elements with limited flexibility for expansion or modifications. In CDE systems, these fixed region-finding pairs typically include a degree or qualifier option but do not allow for the creation of new combinations beyond those predefined in the system. The rationale behind selecting the "region-finding-qualifier" triad approach was twofold. First, it allowed us to perform named entity recognition on the region, finding, and qualifier independently, which greatly facilitated the development of the NLP pipeline. Second, this approach enabled us to deal with a variety of report styles and older reports that might not capture brain features or findings according to modern standards.

To ensure consistency and standardization, we utilized external ontologies and controlled vocabularies for the model attributes. These sources included the Allen Human Brain Atlas (AHBA) [9], Systematized Nomenclature of Medicine (SNOMED) [10], National Cancer Institute Thesaurus (NCIT) [11], Disease Ontology (DOID) [12], Medical Subject Headings (MeSH) [13], Federal Interagency Traumatic Brain Injury Research (FITBIR) CDEs [14], and National Alzheimer’s Coordinating Center (NACC) CDEs [15].

### NLP pipeline and postprocessing

We utilized OpenAI GPT-3.5 model [8] as the primary LLM engine for our NLP pipeline development. No additional fine-tuning was performed, and the standard settings were employed. All calls for gpt-3.5-turbo were executed through the command line, as per the recommended guidelines using default model parameters (temperature 1, top P 1, frequency penalty 0, presence penalty 0). We used python for data extraction and R for data harmonization and Quality Control (QC).

Two approaches were adopted for data extraction from neuropathological reports. The first approach was questionnaire-based, in which the input text was directly mapped to the data model through a series of questions and coded answers. This method facilitated a structured approach to obtaining relevant information from the reports.

The second approach focused on the direct extraction of tissue-finding-qualifier triads from the text. However, this required a harmonization step, as the extracted tissue, finding, and qualifier terms were not directly mapped to our data model. To address this challenge, we performed manual harmonization with the assistance of an embedding-based classification technique, using the text-embedding-ada-002 model from OpenAI [16]. This process involved combining extracted raw terms with terms from controlled vocabularies (e.g., AHBA for regions, the controlled vocabulary for findings and qualifiers), calculating embeddings, and clustering them into groups of 10-20 terms using DBScan. We then manually analyzed each cluster, mapping terms to controlled vocabularies and extending them as necessary. This approach significantly accelerated the mapping process by grouping semantically similar terms, allowing for efficient categorization and mapping of raw extracted terms into a standardized format compatible with our data model, which was based on the aforementioned ontologies and controlled vocabularies.

### Evaluation metrics

To evaluate the quality of data extraction, we compared the results obtained from the automatic extraction pipeline with the manually curated data ("gold standard") for the 65 selected reports. We used the following metrics to assess performance:

- Jaccard Index: Measures the overlap between two sets, calculated as the size of the intersection divided by the size of the union of the sets.
- Sensitivity: The proportion of true positives correctly identified, calculated as true positives divided by the sum of true positives and false negatives.
- Precision: The proportion of positive identifications that are correct, calculated as true positives divided by the sum of true positives and false positives.

For macroscopic and microscopic findings, we assessed the agreement between the lists of brain regions identified manually and those extracted by the automatic pipeline. In cases where brain regions aligned, we compared the associated findings. In cases where both brain regions and findings aligned, we compared the associated qualifiers. For regions with mismatches, we sampled and analyzed regions that were present exclusively in either the manual curation or the pipeline extraction. For neuropathological staging information, we compared the manually curated staging data with the staging information extracted by the automatic pipeline.

By conducting this thorough evaluation, we were able to assess the performance of the extraction pipeline and identify areas for improvement.

## RESULTS

### Data model

#### Overview

In this study, we have developed a comprehensive data model to represent and capture the relevant information from neuropathological reports. Key entities and attributes are shown on the Entity Relationship Diagram (ERD; Figure 1). Full ERD and tabular description of all entities and attributes are provided in Supplementary Information S1 and S2. The primary objective of this data model is to efficiently organize and store the extracted data from the pathology reports, facilitating easy access and analysis for researchers. The data model comprises several key entities, which are interconnected to represent the various aspects of the neuropathological findings. At the core of the data model is the Neuropathological Evaluation entity, which serves as a central hub linking the other entities. Additionally, this entity is connected to the Donor entity, enabling a clear association between the evaluation results and the corresponding donor.

**Figure 1.**
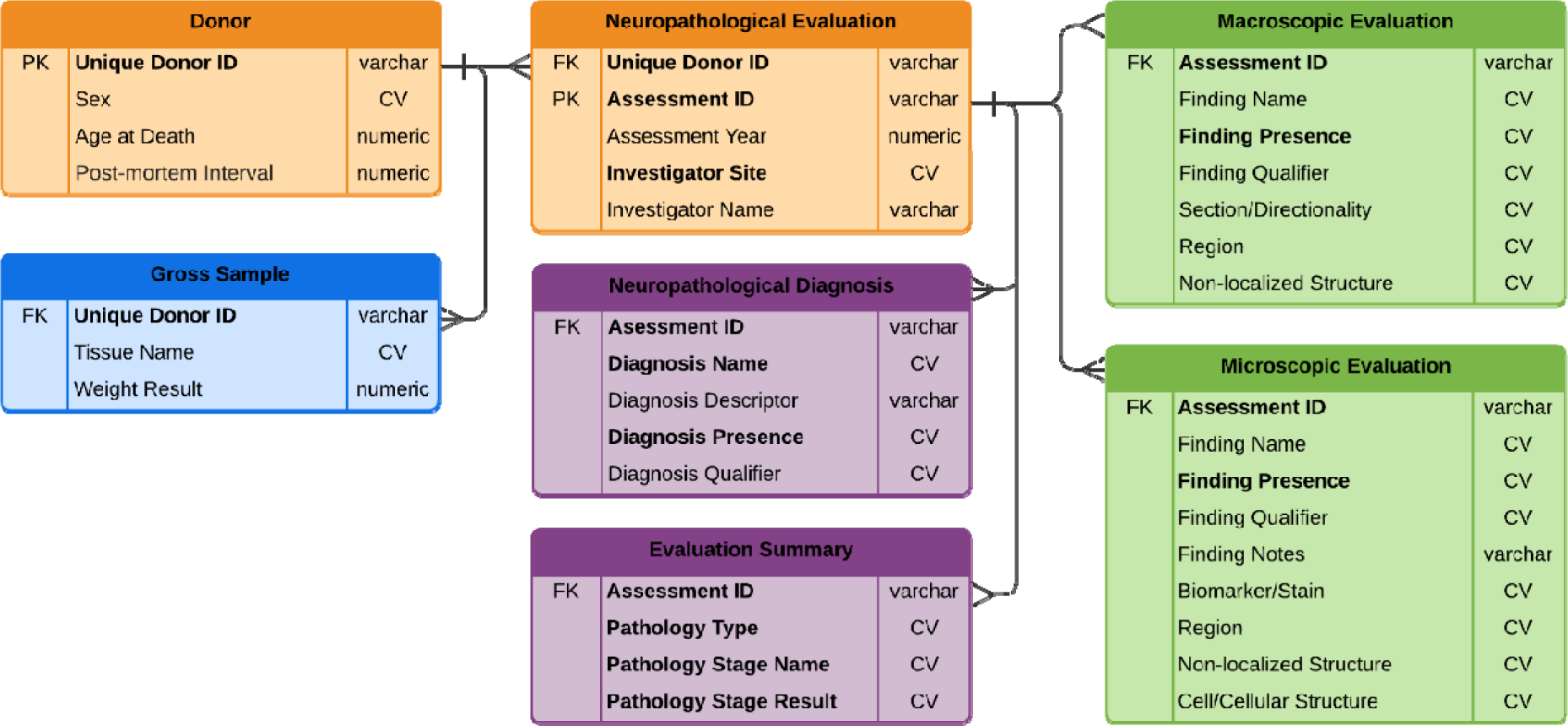
– Entity Relationship Diagram for Neuropathological data model. Key entities and attributes are shown. Relationships between entities follow standard crow foot notation. Color scheme corresponds to conceptual subschemas: orange – donor conceptual subschema, blue – biological specimen conceptual subschema, purple – case diagnosis conceptual subschema, green – pathology case conceptual subschema. Attributes in bold are mandatory. PK – primary key, FK – foreign key, CV – attribute values taken from controlled vocabulary.

The Neuropathological Evaluation entity is further linked to four main sub-entities: Evaluation Summary, Neuropathological Diagnosis, and Macroscopic and Microscopic Evaluations. The Evaluation Summary entity encompasses the staging information (e.g., Braak and Del Tredici stage for PD [17], ABC score according to NIA-AA 2012 consensus guidelines [18], and others [19–22]). The Neuropathological Diagnosis entity lists all neuropathological diagnoses identified in the report, serving as a comprehensive catalog of the patient’s conditions.

The Macroscopic and Microscopic Evaluations entities capture the detailed findings from the gross and microscopic examinations of the brain samples, respectively. These entities store both positive and negative findings, ensuring a complete representation of the pathological landscape. They are essential for understanding the specific pathological abnormalities present in the sample and their implications on the patient’s condition.

#### Data dictionary

To accurately capture the anatomic location of pathology findings, we utilized the AHBA [9] as the foundation for our data model’s representation of brain regions. The AHBA offers comprehensive coverage of brain structures; however, certain adjustments and extensions were necessary to address the specific needs of our study. Firstly, the AHBA does not encompass the vascular system. To address this, we added major arteries to the data dictionary and provided corresponding links to external ontologies such as MeSH [13] or UBERON [23]. Similarly, we included adjacent structures that are not part of the brain, such as the skull and scalp, with appropriate links to external ontologies.

Another challenge we encountered was the presence of hyperspecific and hypospecific regions in the neuropathological reports. Hyperspecific regions, such as the *CA1/CA2 junction* or *calvarial dura*, contain a level of detail absent in the AHBA. Conversely, hypospecific regions, such as the *visual cortex* or *olivary nucleus*, represent groups of brain regions that do not have a corresponding entity in the AHBA. In some cases, the reports used terminology for brain regions that only exist for non-human species, such as the *caudal medullary velum* (rat) or *occipital gyrus* (macaque). To address these issues, we added these regions to our data dictionary and, where possible, provided links to external ontologies and parent regions.

In instances where the report specified a particular part of a brain region, we captured this information using a combination of “Region” and “Section/Directionality”. For example, *posterior occipital cortex* was mapped to a combination of the *occipital cortex* region (AHBA id:3614) and *posterior* directionality. Lastly, we acknowledged that many reports did not associate specific regions with certain findings. For example, reports may mention finding in blood vessels, gray matter, or lesions, without indication of where exactly the finding is located. To accommodate these cases, we included a "Non-localized Structure" attribute in our data model.

In the development of our data model, we aimed to effectively capture and represent macroscopic and microscopic finding names and their associated qualifiers. Generally, finding names encompass descriptions of the observation, such as calcification, atrophy, necrosis, or abnormal coloration, while qualifiers provide optional details regarding severity, quantity, color, shape, and other properties of the findings. We used findings and qualifiers from existing neuropathology CDEs such as those supplied by FITBIR and NACC and extended the dictionary with information from reports. Our initial approach involved separating finding names and qualifiers into basic repeating elements to reduce the number of distinct values in the dictionary and streamline data extraction and harmonization. However, after consultations with the NBB working group, we made certain exceptions. For example, instead of separating *diffuse plaques* into the finding *plaque* and the qualifier *diffuse* we maintained it as a single finding. This decision was made to preserve the specificity and clarity of certain findings.

The final data model comprised 183 macroscopic findings, 416 microscopic findings, and 333 qualifiers. To ensure consistency and interoperability, we mapped the findings to established external ontologies, such as SNOMED and NCIT whenever possible. Specifically, we successfully mapped 78 out of 183 macroscopic findings (42.6%), 97 out of 416 microscopic findings (23.3%), and 38 out of 333 qualifiers (11.4%) to these ontologies.

### Data Extraction Pipeline

#### Overall description

The overall data extraction pipeline (Figure 2) involves six crucial steps. In the first step, the neuropathological reports in pdf and docx file formats are converted into HTML file format, which efficiently preserves the styling information. This conversion allows for easier parsing and extraction of relevant data in subsequent steps. In the second step, the HTML documents are split into distinct sections, such as gross pathology, microscopic findings, diagnosis, and sample information. This splitting is achieved by identifying formatting characteristics (e.g., bold text, larger font size, all caps) that typically denote section headers in the original documents and are preserved in the HTML conversion. For reports containing available tissue information, an additional subcategory titled “tissue” is incorporated. An output table is generated, encompassing report ID, section type, and section content, which serves as a structured representation of the data extracted from the reports.

**Figure 2.**
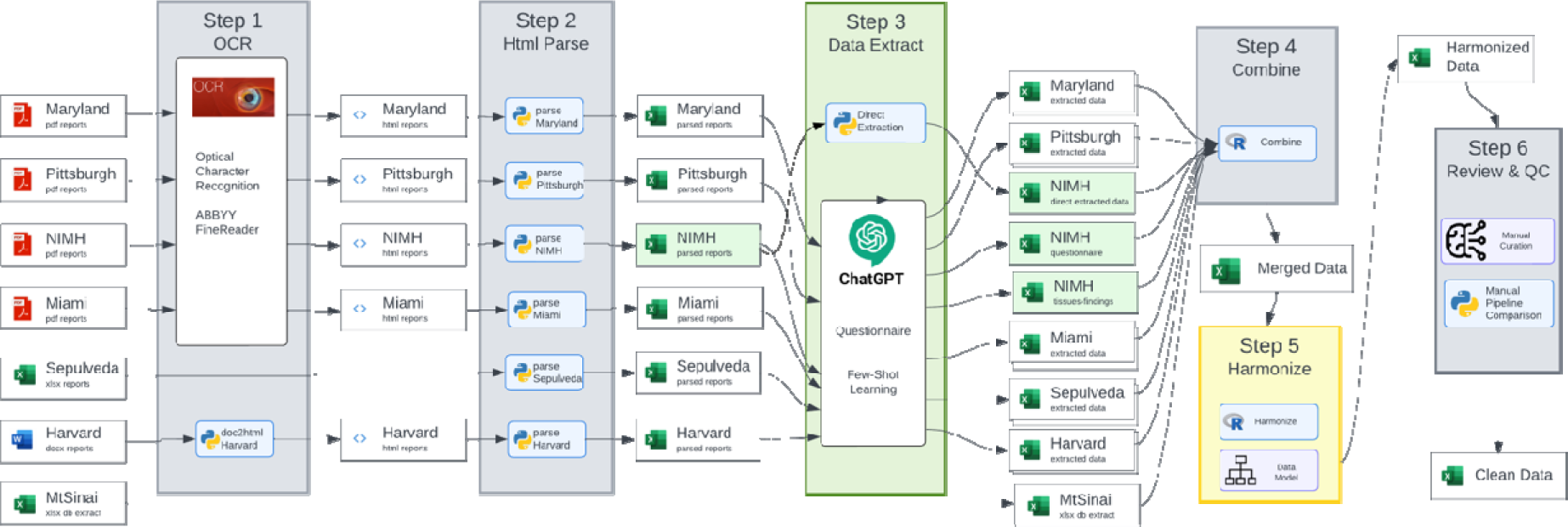
– Schematic representation of data extraction pipeline.

The third step involves feeding the machine-readable data from the output table into the data extraction process. Some information, such as brain weight, donor age, and sex, is already structured and can be effortlessly extracted using pattern matching. To extract most of the other information, we employed two approaches: a questionnaire-based method and a few-shot learning direct approach, both utilizing GPT-based models from OpenAI. In the fourth step, all extracted information is combined and reshaped to create a unified dataset. This dataset then undergoes a harmonization process in step five, where the data is mapped and aligned with the developed data model specifically tailored for neuropathological conditions.

The final step consists of assessing the quality of the extracted data. Values are compared to both the data model and manually curated data to ensure accuracy and consistency across the dataset. Any discrepancies or issues identified during the quality assessment are addressed to refine the data extraction pipeline further.

#### Direct approach for data extraction

The data extraction process adapted an approach conceptually similar to traditional Named Entity Recognition (NER) techniques. Classical NER methods, such as N-gram phonetic search [24], perform optimally when dictionaries are well-defined and comprehensive. However, in our case, we could not rely on the AHBA for region extraction, as it did not encompass all the regions we intended to extract. Furthermore, the dictionary for findings was non-existent, rendering conventional NER tools unsuitable for extracting findings and qualifiers. Consequently, we employed LLMs in the initial step of data extraction to identify all mentioned brain regions. To improve sensitivity, region extraction was executed twice, and tissue lists from both runs were consolidated. Subsequently, for each mentioned tissue, LLMs extracted associated findings and qualifiers. We guided the model using a few-shot learning approach, providing request and response examples to assist in handling complex or ambiguous cases. Examples were selected based on analysis of cases where initial extraction failed, with 4-6 examples typically included per prompt. Prompts were tailored slightly for different NBB sites.

Initially, the davinci-03 model was employed, which was not specifically fine-tuned for user reque ts. In the final version, we used the gpt-3.5-turbo model. Since gpt-3.5-turbo was trained to respond to direct user requests, we supplemented the prompts with explicit instructions regarding the desired output format. Examples and the overall protocol can be found in Figure 3.

**Figure 3.**
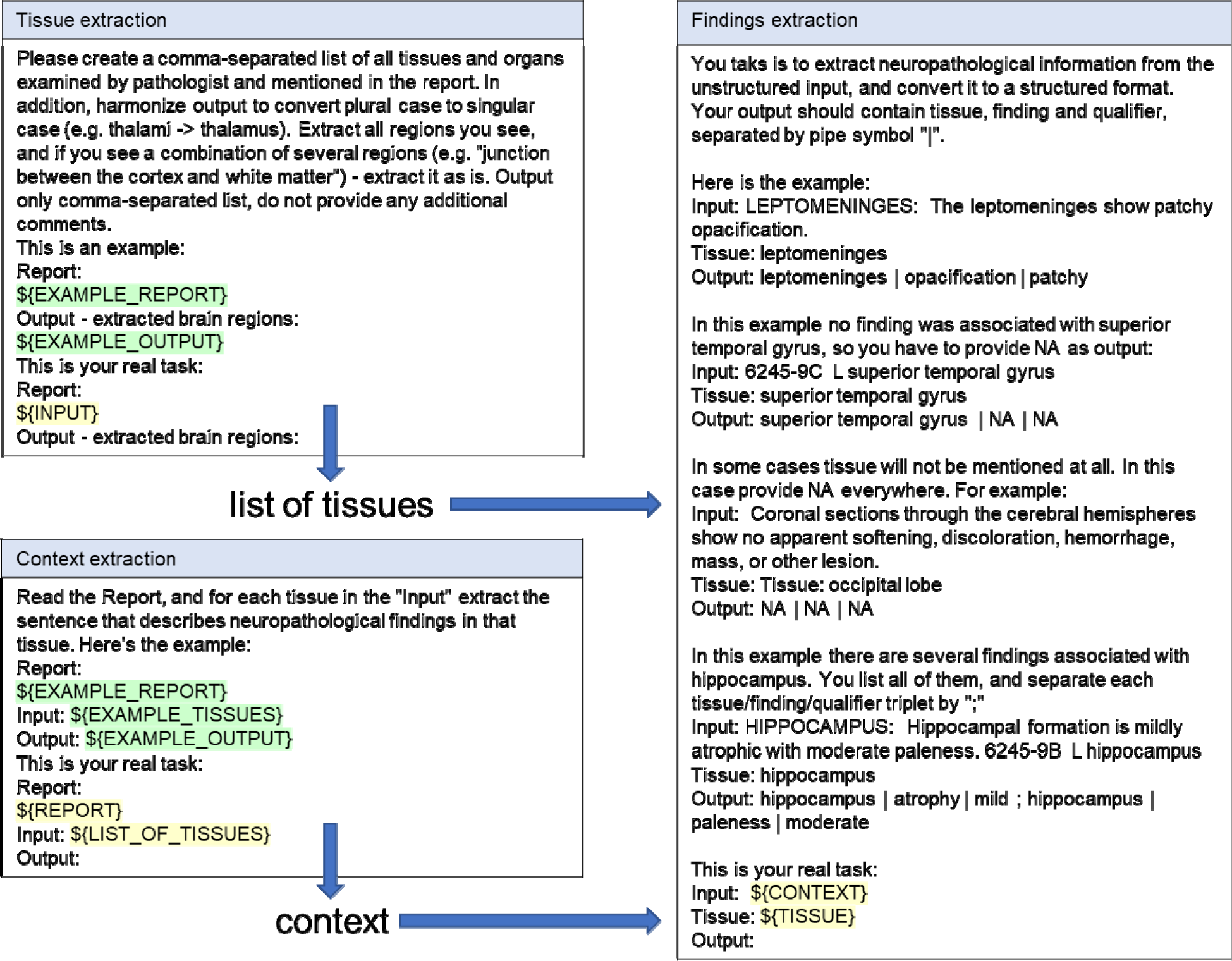
Workflow for direct data extraction. The $ designates text that is dynamically substituted by the data from reports, or results obtained on previous steps.

While the direct approach effectively captured information present in the reports, it necessitated subsequent harmonization, as the LLM was not cognizant of specific dictionaries employed in later stages. Moreover, the model did not consistently differentiate between finding names and qualifiers, resulting in discrepancies such as Finding Name *size decreased* versus the combination of Finding Name *size* and Finding Qualifier *decreased*. Therefore, a crucial harmonization step was essential to render the extracted information useful and consistent. Nevertheless, the developed few-shot learning approach successfully navigated ambiguous information and the absence of dictionaries, yielding semi-structured raw output that could be harmonized downstream.

#### Questionnaire-based approach for data extraction

The questionnaire-based approach emulates data abstraction by filling electronic forms. In this method, a series of questions are posed to the text, with coded answers provided. The goal is not to extract every piece of data but to focus on what is most important, as defined by established domain-specific questionnaires. These questionnaires, developed by experts in the field, prioritize most clinically and research-relevant information for specific diseases. For example, the NACC questionnaire focuses on findings crucial for establishing diagnoses of AD and PD. We employed a similar approach where, instead of a human operator, it was the LLM that answered the questions. In brief, the LLM was provided with instructions to answer the questions, a context (specific section of the report), and the questions themselves. These instructions or questions cold be direct (e.g., “Provide the whole brain weight, in grams.”) or dictionary-based (e.g., “What is the severity of cerebral cortex atrophy?” with coded answer options that were taken from NACC or FITBIR questionnaires). Answers to dictionary-based questions were mapped directly to the data model, so no additional harmonization was required.

Assessing data extraction quality for data extracted by the questionnaire revealed several failure modes. First, the LLM was not able to discriminate between cases where the region was not mentioned and cases where no abnormalities were found. Moreover, the LLM tended to deviate from direct answers and make conclusions that were close but not exactly answering the questions. For example, when asked about the Braak & Del Tredici stage of PD [17], the model sometimes attempted to interpret the extent of findings and give stage assessments, rather than reporting that the stage was not mentioned (Figure 4). To address these issues, we modified the flow of the questionnaire and added intermediate questions such as “Is X mentioned in the report? (0- No, 1- Yes)”, “Is there evidence of X? (0- No, 1- Yes)”, and only then “What is the severity of X?”. This approach enabled us to distinguish more precisely between present and absent findings and reduce hallucinations.

**Figure 4.**
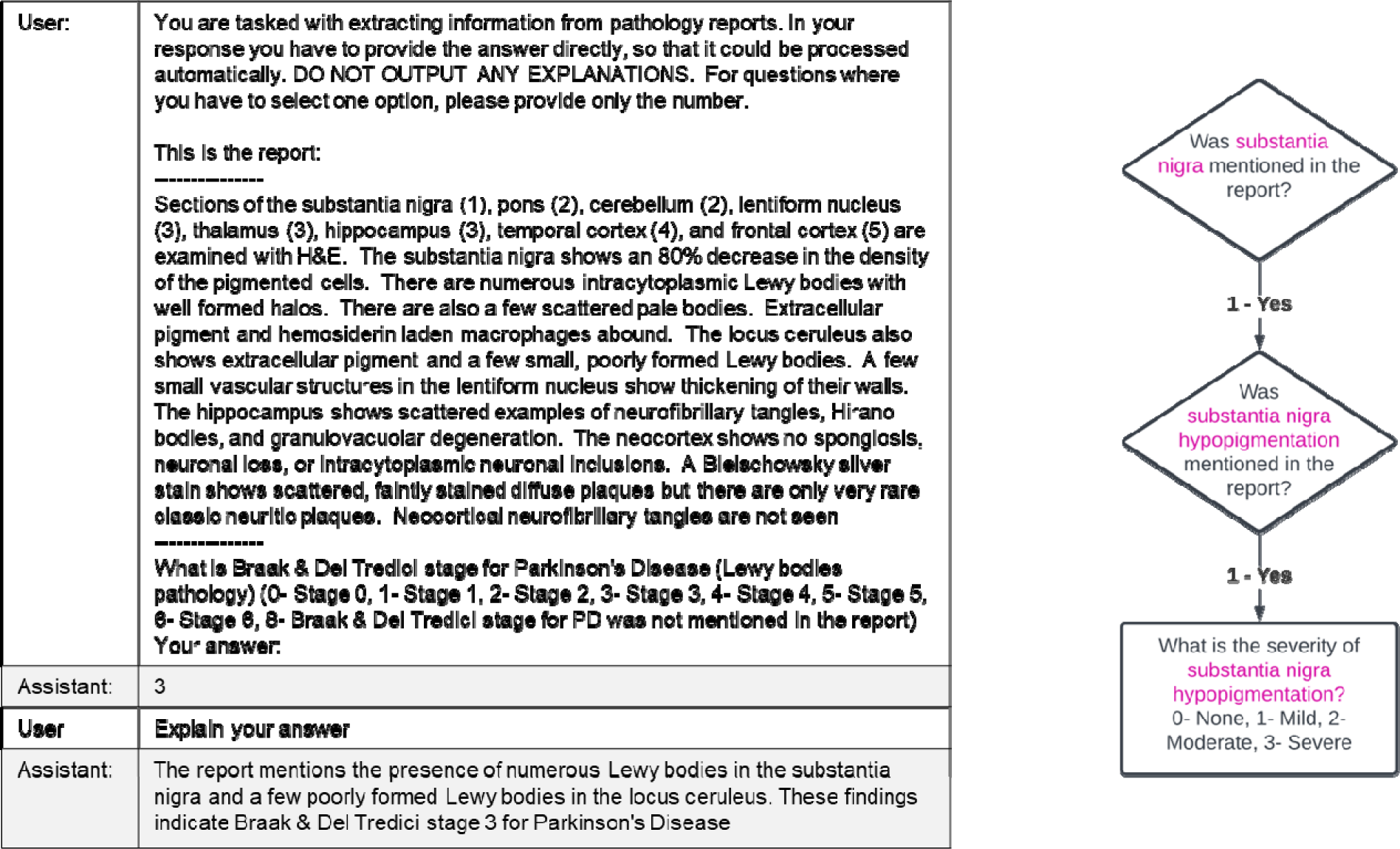
An example of a typical questionnaire prompt with responses and follow-up. Flowchart of the questions aimed to discriminate between absence of information, negative findings, and positive findings. The follow-up question ’Explain your answer’ was used only during manual debugging to understand the LLM’s reasoning and to develop examples for few-shot learning. It was not included in the final automated pipeline.

### Quality Assessment

#### Overall quality assessment results

To assess the quality of data extraction, we compared the data extracted from 65 reports by the pipeline to the data extracted through manual curation. For macroscopic and microscopic findings, we evaluated the agreement between the lists of brain regions identified manually and those extracted by the automatic pipeline. In cases where findings intersected, we compared the associated qualifiers. For regions with mismatches, we sampled and analyzed regions that were exclusively present in either the manual curation or pipeline extraction. For neuropathological staging information, we compared the manually curated staging data with the staging data extracted by the automatic pipeline. Overall metrics are reported in Table 1.

**Table 1.** Overall QC metrics for Macroscopic and Microscopic evaluation (comparison of sets of regions) and Evaluation summary (comparison of staging information). C and P denote information extracted by Curation and Pipeline respectively. C∩P denotes records intersection of sets of records, C[P denotes union of sets.

|  | Jaccard Index<br>(accuracy) | Sensitivity | Precision |
| --- | --- | --- | --- |
| | $(C \cap P) / (C \cup P)$ | $(C \cap P) / C$ | $(C \cap P) / P$ |
| Macroscopic evaluation | 64.6% | 83.6% | 73.9% |
| Microscopic evaluation | 55.9% | 76.5% | 67.5% |
| Evaluation summary | 64.6% | 71.9% | 86.4% |

To further characterize problems with data extraction, we sampled up to 10 mismatches of both types (data present in manual curation only, data present in pipeline extraction only) for every site for regions in macroscopic evaluations. Mismatches were categorized based on the type of problem: issues with pipeline data extraction, issues with manual curation, or issues with harmonization. Then, types of issues were identified (Table 2).

**Table 2.** Breakdown of comparison between manually extracted data and data extracted by pipeline by category for macroscopic evaluation. Error percentage is estimated from sampling of up to 20 mismatching records per site (10 records with curation only, 10 records with pipeline only region).

| Category | Description | Estimated % |
| --- | --- | --- |
| Match | Results of curation matches results of the pipeline | 64.6% |
| Pipeline issues | Any issues due to incorrect or imprecise data extraction by pipeline. This includes the following sub-categories: | 9.9% |
| Pipeline errors | Information was extracted by pipeline incorrectly | 3.7% |
| Pipeline recovery | Brain regions are present in report but were not extracted by the pipeline | 2.5% |
| Pipeline precision | Regions were extracted by pipeline with insufficient precision | 1.9% |
| Pipeline missing/normal | Findings were extracted by pipeline as normal, but were missing in report or vice versa | 1.9% |
| Curation issues | Any issues due to incorrect or imprecise data extraction by curators. This includes the following sub-categories: | 9.6% |
| Curation recovery | Brain regions are present in report but were not extracted by curators | 5.2% |
| Curation precision | Regions were extracted by curators with insufficient precision | 4.4% |
| Harmonization issues | Mismatch due to information being extracted in different ways, or mapped to different terms | 15.8% |

Issues with the pipeline data extraction comprised 9.9% of the total amount of problems. In 9.6% of cases, information extracted by the pipeline was more accurate than information produced by curators. Additionally, 15.8% of mismatches were due to issues with harmonization, where both curation and pipeline approaches produced results that could be considered valid – the same data was interpreted in different ways. This analysis suggests that the total accuracy of data extraction for macroscopic findings surpasses 74.2% (the number of matching records plus the number of issues due to curation problems).

### Analysis of common pipeline extraction errors

#### Incorrect data extraction

We identified four types of pipeline errors. The most frequent cases involved incorrect data extraction (3.7%). In many instances, these errors resulted from the incorrect extraction of context. For example, the LLM accurately identifies “cerebral hemisphere” as one of the brain regions mentioned in the following extract:

*“Coronal sections of the left hemisphere at the anterior frontal, striatal, and lentiform-thalamic- substantia nigra levels, and the midpons-cerebellum are examined. There is no softening, discoloration, hemorrhage, mass, or other lesion. Moderate cortical atrophy is seen in the frontal and temporal lobes.”*

The model then proceeds to extract the following context: *Coronal sections of the left hemisphere at the anterior frontal, striatal, and lentiform-thalamic-substantia nigra levels, and the midpons-cerebellum are examined. Moderate cortical atrophy is seen in the frontal and temporal lobes.*”, which is then converted into a region, finding, and qualifier combination of “***cerebral hemisphere***, *atrophy*, *moderate*”. This is less precise than the “***frontal cortex***, *atrophy*, *moderate*” extracted by curators.

#### Extraction formatting errors

In 2.5% of cases, the pipeline did not extract region information. Many of these errors occurred when the tissue was correctly identified, but the pipeline had trouble producing results in the desired format. For example, “*The lateral cerebral ventricle is normal in size and shape*” was incorrectly converted to “*lateral cerebral ventricle | **normal size; normal shape** | NA*”. In the correct format, region/finding/qualifier triplets should be separated by semicolons, and the correct output should look like “*lateral cerebral ventricle | normal size | NA; lateral cerebral ventricle | normal shape | NA*”.

#### Imprecise data extraction

Imprecise extraction accounted for 1.9% of the pipeline extraction problems. Typical examples included extraction of “cerebral cortex” without specifying the pathology location in more detail. In many cases, the pipeline extracted both general regions (*cerebral cortex*) and specific regions (e.g., *frontal cortex*). Additionally, some questions from the questionnaire asked about the presence of pathology in general regions and were therefore mapped to these general areas. Moreover, the coded list of answers forced the LLM to perform qualifier mapping. For instance, *“minimal”* was typically mapped to *“mild”*, whereas “*mild to moderate*” could be mapped to either *“mild”* or *“moderate”* without a systematic approach to the mapping process. Similarly, pathological evaluations often contained ranges of stages (e.g., Braak II-III). The LLM randomly collapsed the range to one of the stages.

#### Confusion between absence of abnormalities and missing information

Lastly, 1.9% of pipeline extraction problems stemmed from confusion between findings that were not reported and cases where no abnormalities were found in specific regions. The LLM would easily become confused when the report contained a full list of sections and tissues that were analyzed, assuming that if a tissue was reported but no abnormality was explicitly mentioned, the tissue was normal. Initially, the percentage of these errors was much higher, so we had to remove the list of regions at the preprocessing stage whenever it was possible to identify such a list using style markers or headers.

#### Harmonization issues and problems with manual curation

Harmonization issues accounted for the largest portion (15.8%) of mismatches between manually curated data and data extracted by the pipeline. Common examples of these issues include discrepancies in naming general regions. For instance, “*cerebrum*” is often interchanged with “*cerebral hemispheres*” by both the pipeline and manual curation.

Another example illustrating the differing approaches between curators and the pipeline can be seen in cases where a list of tissues is reported. For example, a report might contain the following information: *“**Neostriatum**: (caudate nucleus, putamen, and nucleus accumbens): Unremarkable.”* Curators interpreted this as *“neostriatum, unremarkable”* whereas the pipeline extracted all four mentioned tissues separately (*neostriatum*, *caudate nucleus*, *putamen*, and *nucleus accumbens*) and reported all of them as unremarkable.

In some cases, harmonization problems arose from instances where a finding or region could be represented in different ways, such as “*junction between cortex and white matter*” versus a combination of “*cortex*” and “*white matter*”, or “*tonsillar herniation*” versus a combination of “*cerebellar tonsil*” and “*herniation*”.

Lastly, a significant portion of mismatches (9.6%) could be attributed to either imprecise extraction by curators (5.2%) or information not being extracted at all (4.4%). These issues are more prevalent for sites that provide information-rich reports (e.g., Harvard and Miami), as the sheer amount of information in these reports makes manual curation more prone to errors.

### Dataset characteristics

The final version of the dataset extracted from 822 reports on donors with clinical diagnosis of Parkinson’s disease consisted of 19,000 macroscopic and 44,000 microscopic observations. 35% of macroscopic, and 46% of microscopic observations were regarded as abnormal, the remaining could be considered “normal” as the tissues were examined, but no abnormalities were found. 1735 neuropathological diagnosis records were extracted from reports. Lewy body disease was most frequent among the donors; together with Parkinson’s disease it accounted for 689 cases (90.1%) of the 764 reports where information about neuropathological diagnoses was present. Among other frequent diagnosis were Cerebral infarction (234 cases), Cerebral autosomal dominant arteriopathy (230), Frontotemporal dementia (140 cases), see Table 3.

**Table 3.** Number of reports with neuropathological diagnosis (only diagnosis with prevalence > 5% are shown)

| Diagnosis name | Number of reports (%) |
| --- | --- |
| Lewy body disease | 522 (68.3) |
| Cerebral infarction | 233 (30.5) |
| Cerebral autosomal dominant arteriopathy | 230 (30.1) |
| Parkinson's disease | 227 (29.7) |
| Neurofibrillary degeneration | 133 (17.4) |
| Cerebrovascular disease | 100 (13.1) |
| Frontotemporal dementia | 60 (7.9) |
| Alzheimer's disease | 49 (6.4) |
| Total number of neuropathological diagnoses | 764 (100) |

Interestingly, 10% donors did not have Lewy body disease or Parkinson’s disease neuropathological diagnoses, despite having clinical diagnoses. Most frequent neuropathological diagnoses for such patients were Cerebral autosomal dominant arteriopathy (50 cases), Cerebral infarction (50 cases) and Frontotemporal dementia (26 cases).

Most frequent microscopic and macroscopic findings are presented in Table 4. Not all reports contained results of macroscopic or microscopic examination. Hypopigmentation of substantia nigra was present in 59.5% of 773 reports that contained data on macroscopic data. Microscopically, these findings were frequently correlated with neuronal loss and Lewy bodies. Among other macroscopic findings were atrophy (in frontal lobe, or without further specification), pigmentation of substantia nigra, and increased size of lateral ventricle. Microscopically, neuronal loss and Lewy bodies were observed in substantia nigra and more specifically in pars compacta, and locus coeruleus, neurofibrillary degeneration – in entorhinal area and hippocampus.

**Table 4.** 10 most frequent microscopic and macroscopic findings extracted from reports.

| Finding | Brain region | Number of reports (%) |
| --- | --- | --- |
| Macroscopic findings |  | 773 (100) |
| Hypopigmentation | substantia nigra | 485 (59.5) |
| Hypopigmentation | locus coeruleus | 285 (35.0) |
| Atrophy | frontal lobe | 237 (29.1) |
| Atrophy | cerebral cortex | 229 (28.1) |
| Pigmentation | substantia nigra | 203 (24.9) |
| Increased size | lateral ventricle | 153 (18.8) |
| Atherosclerosis | cerebrum | 124 (15.2) |
| Atrophy | temporal lobe | 123 (15.1) |
| Atherosclerosis | circle of Willis | 109 (13.4) |
| Microscopic findings |  | 815 (100) |
| Neuronal loss | substantia nigra | 434 (53.3) |
| Lewy bodies | substantia nigra | 403 (49.4) |
| Neurofibrillary degeneration | entorhinal area | 295 (36.2) |
| Lewy bodies | locus coeruleus | 267 (32.8) |
| Neuronal loss | locus coeruleus | 231 (28.3) |
| Neurofibrillary degeneration | hippocampus | 206 (25.3) |
| Neuronal loss | pars compacta | 197 (24.2) |
| Lewy bodies | pars compacta | 168 (20.6) |
| Gliosis | substantia nigra | 164 (20.1) |
| Pigmentation | pars compacta | 158 (19.4) |

## DISCUSSION

In this study, we explored the capabilities of LLMs, such as GPT-based models, for data extraction from neuropathological reports. The first step for successful data extraction involves building a flexible data model that can accommodate a variety of data while facilitating extraction and subsequent harmonization. We achieved this by centering our model on regions, findings, and qualifiers, each of which can be varied independently. This approach deviates from the approach used to build CDEs, in which the central conceptual entity consists of predefined sets of regions and findings. While the CDE approach is more suitable for standardizing crucial information in electronic form, it is less flexible and less suitable for automated data extraction. Moreover, different electronic systems may contain sets of CDEs which are not compatible. By deconstructing CDEs into triads of regions, findings, and qualifiers, data becomes more flexible and can be easily converted, as demonstrated in our approach where we merged data from electronic system used to capture information in Mount Sinai with data extracted from pathology reports from other sites.

This study has demonstrated the significant potential of LLMs in structuring unstructured neuropathological reports. LLMs can handle ambiguous information and the absence of predefined dictionaries, which is essential when dealing with diverse and complex data. The reasoning capabilities of LLMs make it possible to extract complicated relationships and infer information that would be unreachable for standard methods. In certain cases where the amount of information is overwhelming, LLMs outperform manual data extraction and retrieve information more reliably.

Despite the benefits of LLMs, there are limitations to their application in data extraction from neuropathological reports. For instance, LLMs may struggle to differentiate between cases where a region is not mentioned and cases where no abnormalities were observed. Additionally, LLMs may inaccurately extract context or become confused when dealing with complex report structures. These limitations can lead to discrepancies between manually curated data and data extracted by the pipeline. Furthermore, LLMs may occasionally deviate from direct answers or make conclusions that are close but not exactly answering the questions.

The present study serves as a pilot effort, focusing on donors with PD, which represents less than 5% of the total number of pathology reports in the NBB. Although a significant portion of donors with PD have other comorbidities (e.g., AD and Cerebrovascular Disease), extending the pipeline to work with other patients would necessitate modifications to both the data model and the extraction workflow. For instance, entities describing brain tumors or specific findings related to traumatic brain injury could be incorporated into the model. The data dictionary could be expanded to encompass findings more characteristic of other pathologies, and corresponding evaluation staging information should be included. Moreover, certain sites (Pittsburgh and NIMH) were sparsely represented in the PD datasets we examined, and integrating additional reports from these sites might require modifications to the preprocessing algorithm. Additionally, the NACC questionnaire used in the data extraction pipeline should be extended with questions relevant to all types of pathologies present in the broader population of reports.

While the current data extraction approach achieved 74.2% accuracy and was comparable with manual curation results, we suggest several avenues for further improvement and scaling up. The LLM used in this study, GPT-3.5, has been superseded by a more powerful model, GPT-4 [25]. Another approach may involve fine-tuning the LLM to better understand the specific domain of neuropathology and further adapting it to handle complex and diverse data formats.

A significant portion of time was devoted to manual harmonization of data and mapping from LLM output to data dictionaries. For some attributes such as brain regions and qualifiers, we expect the harmonization efforts to scale sub-linearly, as the current set of reports already covered a significant variety of values. Other attributes, such as finding names, might still require substantial efforts to harmonize, as the data is dependent on the type of pathology. LLMs and related NLP capabilities, such as using embeddings for mapping between information provided by LLMs and predefined ontologies, might be employed to expedite the harmonization process.

The resulting dataset contains a substantial volume of information on macroscopic and microscopic pathology of Parkinson’s disease. Due to the presence of significant amount of other pathologies such as cerebral infarction, Frontotemporal dementia and Alzheimer’s disease, the pipeline and data model could be easily extended to other neuropathological conditions.

## CONCLUSIONS

In this study, we have developed a data model and data extraction pipeline that leverages LLMs to structure unstructured neuropathological reports from the NBB, specifically focusing on PD donors. To our knowledge, this is the first attempt to automatically standardize neuropathological information at this scale. The pipeline and data model can be repurposed and extended to accommodate other pathological conditions, making it a versatile tool for researchers. Furthermore, the collected data has the potential to serve as a valuable resource for PD researchers, bridging the gap between clinical information and genetic data, and thereby facilitating a more comprehensive understanding of the disease.

## ACKNOWLEDGEMENTS

Data dictionary developed in this study was built using the controlled access datasets distributed from the DOD- and NIH-supported Federal Interagency Traumatic Brain Injury Research (FITBIR) Informatics Systems. FITBIR is a collaborative biomedical informatics system created by the Department of Defense and the National Institutes of Health to provide a national resource to support and accelerate research in TBI.

Brain pathology reports were provided by NIH NeuroBioBank, including Mount Sinai Neurobiobank, University of Miami Brain Endowment Bank, University of Maryland Brain and Tissue Bank, Harvard Brain Tissue Resource Center, The Human Brain and Spinal Fluid Resource Center, Brain Tissue Donation Program at the University of Pittsburgh, National Institute of Mental Health’s Human Brain Collection Core.

Authors would like to thank Dr. Harry Haroutunian, Dr. William Scott, Dr. Derek Oakley, Dr. Pavan Auluck, Dr. Stefano Marenco, and NIH BioBank directors for their advice in development of the data model.

## FUNDING

The work described herein was created with funding by NINDS (Contract: HHSN316201200128W- 75N95022F00001)

## CODE AVAILABILITY

The code used in this study can be made available upon reasonable request to the corresponding author.

## SUPPORTING INFORMATION

S1 – Neurological Data Model – entities and attributes specification (in Excel file format) S2 – Entity Relationship Diagram for Neurological Data Model

S3 – Composition of the PD reports subset for manual curation (breakdown by site) S4 – General statistics on manually curated subset of PD reports

## Notes

### Competing Interest Statement

The authors have declared no competing interest.

### Summary of Updates

added "Dataset Characteristics" section to the results; Methods section expanded

