## Supplementary figures and images for "Unpacking Unstructured Data: A Pilot Study on Extracting Insights from Neuropathological Reports of Parkinson’s Disease Patients using Large Language Models"

### Supplementary Data 2

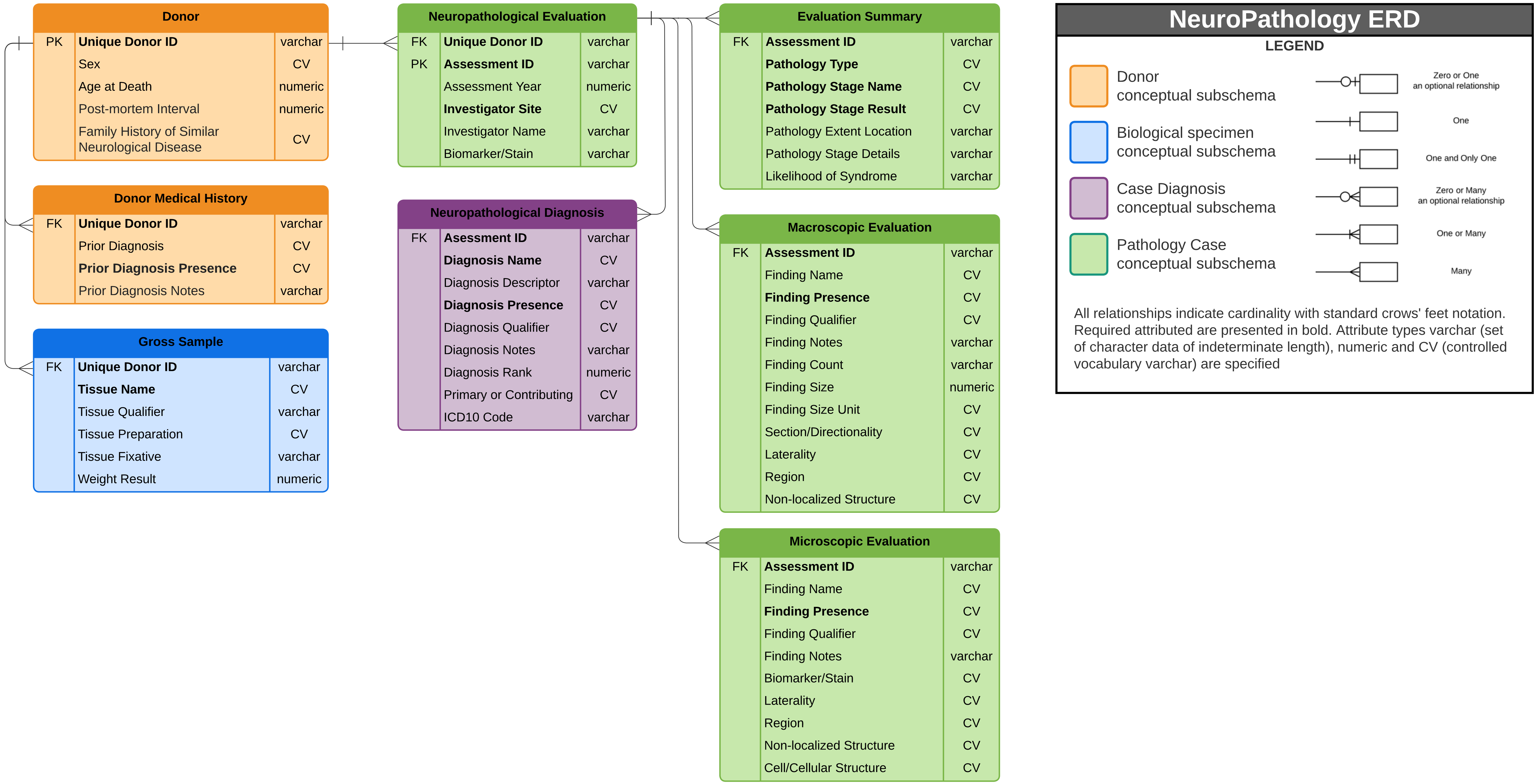
