## Supplementary Table 3 for "Unpacking Unstructured Data: A Pilot Study on Extracting Insights from Neuropathological Reports of Parkinson’s Disease Patients using Large Language Models"

Supplementary Table S3 – Composition of the PD reports subset for manual curation (breakdown by site)

| NBB site | Number of reports | Number of formats | File type |
| --- | --- | --- | --- |
| Maryland | 12 | 4 | pdf |
| MtSinai* | 3 | 1 | pdf, excel |
| Harvard | 16 | 2 | doc |
| Sepulveda | 16 | 3 | pdf |
| Pittsburgh** | 1 | 1 | pdf |
| NIMH** | 5 | 2 | pdf |
| Miami | 12 | 2 | pdf |

* Majority of MtSinai reports were already digitized and were provided in excel format. Only 3 non-digitized reports on PD patients were provided by NBB.

** PD donors were underrepresented in Pittsburgh and NIMH biorepositories compared to other sites
