## Supplementary Table 4 for "Unpacking Unstructured Data: A Pilot Study on Extracting Insights from Neuropathological Reports of Parkinson’s Disease Patients using Large Language Models"

Supplementary Table S4 – General statistics on manually curated subset of PD reports

| Parameter | Value |
| --- | --- |
| Total number of reports | 65 |
| Number of sites | 7 |
| Number of different formats | 15 |
| Number of reports that has did not have text layer and required OCR (%) | 16 (24.6%) |
| Average size of report, pages | 3.875 |
| Average length (in words) of reports sections: |  |
| Microscopic findings | 467 |
| Macroscopic findings | 220 |
| Diagnosis | 50.2 |
| Fraction of reports (in %) that have sections: |  |
| Microscopic findings | 93.5 % |
| Macroscopic findings | 90.3 % |
| Diagnosis | 90.3 % |
